# Elevated pyrimidine dimer formation at distinct genomic bases underlie promoter mutation hotspots in UV-exposed cancers

**DOI:** 10.1101/404434

**Authors:** Kerryn Elliott, Martin Boström, Stefan Filges, Markus Lindberg, Jimmy Van den Eynden, Anders Ståhlberg, Anders R. Clausen, Erik Larsson

**Affiliations:** Department of Medical Biochemistry and Cell Biology, Institute of Biomedicine, The Sahlgrenska Academy, University of Gothenburg, SE-405 30 Gothenburg, Sweden; Sahlgrenska Cancer Center, Department of Pathology and Genetics, Institute of Biomedicine, Sahlgrenska Academy at University of Gothenburg, Medicinaregatan 1F, 413 90 Gothenburg, Sweden; Wallenberg Centre for Molecular and Translational Medicine, University of Gothenburg, Sweden; Department of Clinical Pathology and Genetics, Sahlgrenska University Hospital, 413 45 Gothenburg, Sweden

## Abstract

Sequencing of whole cancer genomes has revealed an abundance of recurrent mutations in gene-regulatory promoter regions, in particular in melanoma where strong mutation hotspots are observed adjacent to ETS-family transcription factor (TF) binding sites. While sometimes interpreted as functional driver events, these mutations have also been suggested to be due to locally inhibited DNA repair or, alternatively, locally increased propensity for UV damage. Here, we provide evidence that base-specific elevations in the efficacy of UV lesion formation underlie these mutations. First, we find that low-dose UV light induces mutations preferably at a known ETS promoter hotspot in cultured cells even in the absence of global or transcription-coupled nucleotide excision repair (NER), ruling out inhibited repair. Further, by genome-wide mapping of cyclobutane pyrimidine dimers (CPDs) shortly after UV exposure and thus before DNA repair, we find that ETS-related mutation hotspots exhibit a strong base-specific increase in CPD formation frequency. Analysis of a large whole genome cohort illustrates the widespread contribution of this effect to recurrent mutations in melanoma. While inhibited NER underlies a general increase in somatic mutation burden in regulatory regions, we conclude that the most recurrently mutated individual DNA bases arise instead due to locally favorable conditions for UV damage formation, thus explaining a key phenomenon in whole-genome cancer analyses.

## INTRODUCTION

Whole genome analysis of cancer genomes has the potential to reveal non-coding somatic mutations that drive tumor development, but it remains a major challenge to separate these events from non-functional passengers. The main principle for identifying drivers is recurrence across independent tumors, suggestive of positive selection, which led to the recent identification of frequent oncogenic mutations in the promoter of telomere reverse transcriptase (*TERT*) that can activate its transcription ^1,2^. However, mutation rates vary across the genome, and local elevations may give rise to “false” recurrent events that can be misinterpreted as signals of positive selection. While known covariates of mutation rate, such as replication timing and local trinucleotide context, can be accounted for to improve interpretation ^3^, the non-coding genome may be particularly challenging. Mutational fidelity may be generally reduced in this vast and relatively unexplored space, as indicated by the presence of mechanisms directing DNA repair specifically to exonic regions^4^, and yet-unexplained mutational phenomena may be at play.

Indeed, recent studies have described a remarkable abundance of recurrent promoter mutations in melanoma and other skin cancers, often noted to overlap with sequences matching the recognition element of ETS family transcription factors (TFs) ^5,6,7,8,9,10,11^. Strikingly, a large proportion of frequently recurring promoter mutations in melanoma occur at distinct cytosines one or two bases upstream of TTCCG elements bound by ETS factors as indicated by ChIP-seq, within a few hundred bases upstream of a transcription start site ^12^. While often interpreted as driver events, we recently showed that these sites exhibit highly elevated vulnerability to UV mutagenesis, as evidenced by their rapid induction following low-dose UV light exposure in cultured cells ^12^. The effect has sometimes been attributed to locally impaired nucleotide excision repair (NER) caused by binding of ETS TFs ^11,13,14^. However, our analysis of skin tumors lacking global NER (*XPC* -/-) contradicted this model ^12^ and the mechanism remains unclear. An understanding of this phenomenon, which may underlie a large part of all non-coding recurrent events in human tumors beyond *TERT* ^5,7,11^, would resolve a key question that continues to confound whole cancer genome analyses.

Here, through analysis of 221 whole tumor genomes, we first demonstrate the widespread impact of TTCCG-related mutagenesis on the mutational landscape of melanoma. Moreover, through UV exposure of a panel of repair-deficient human cell lines, we rule out inhibited DNA repair as an important mechanistic basis for ETS-related recurrent promoter mutations in UV-related cancers. Finally, we generate the highest resolution map of UV-induced cyclobutane pyrimidine dimers (CPDs) in the human genome to date, which provides clear evidence that ETS-related promoter hotspots instead arise due to an exceptional local elevation in the efficacy of UV lesion formation at specific genomic bases.

## RESULTS

### Widespread contribution from TTCCG-related sites to recurrent non-coding mutations in 221 melanoma whole genomes

To assess the impact of TTCCG-related mutagenesis on the landscape of recurrent mutations in melanoma in a more sensitive way than previously possible, we assembled a cohort of 221 melanomas characterized by whole genome sequencing by TCGA and ICGC ^15,16^. These heavily mutated tumors averaged 110k somatic single nucleotide variants (SNVs) per sample, expectedly dominated by C>T transitions and a mutational signature characteristic of mutagenesis by UV light through formation of pyrimidine dimers (**Fig. S1**).

Notably, despite the genome-wide scope, nearly all highly recurrent mutations were found near annotated transcription start sites (TSSs) (**Fig. 1a**). For example, of the 22 most recurrent individual bases (mutated in ≥18 patients), four were known drivers (*BRAF, NRAS* or *TERT* promoter mutations) while the rest were at most 524 bp away from a known TSS. Further, the vast majority of highly recurrent promoter sites were found in conjunction with TTCCG sequences (**Fig. 1a-b**), indicating a widespread influence from ETS elements to the mutational landscape of melanoma.

**Figure 1.**
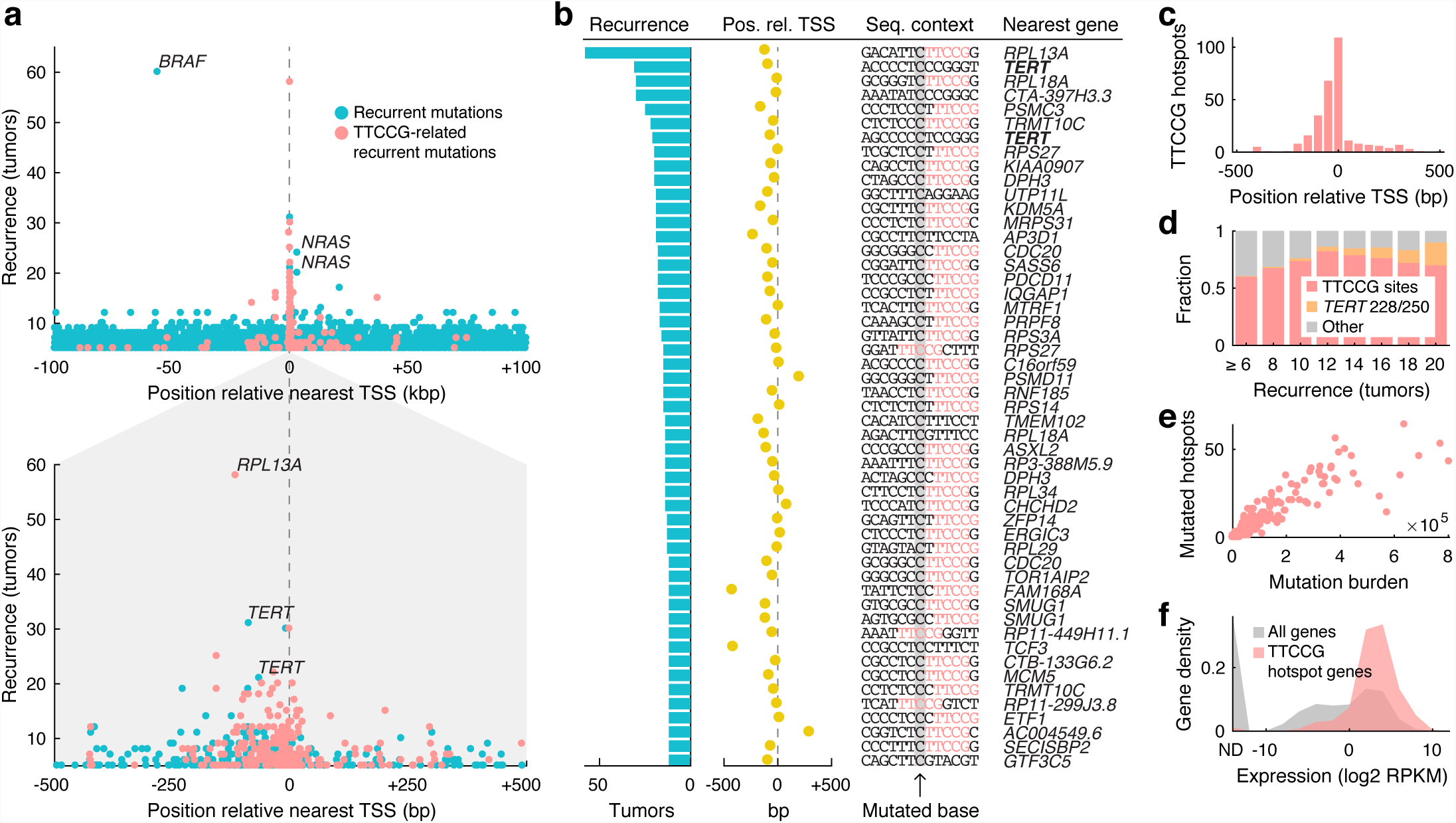
Widespread contribution from TTCCG-related sites to recurrent non-coding mutations in 221 whole melanoma genomes. (**a**) Highly recurrent somatic mutations (individual genomic bases) aggregate near annotated transcription start sites (TSS) and typically colocalize with TTCCG elements. Recurrent sites having a TTCCG element within a +/-10 bp context on the mutated (pyrimidine) strand are indicated (red). Bottom panel: +/-500 bp close-up around the TSS. (**b**) Top 51 recurrent promoter sites (+/-500 bp), all mutated in ≥12/221 tumors (>5%). Degree of recurrence, position relative to TSS, sequence context with TTCCG highlighted in red, and nearest gene are indicated. (**c**) Positional distribution of TTCCG-related mutation hotspots near TSSs, based on 291 promoter sites recurrent in ≥5 tumors. (**d**) Proportion of recurrent promoter mutations (+/-500 bp) that are TTCCG-related (red), *TERT* activating mutations (C228T/C250T; orange) or other (gray), as a function of recurrence. (**e**) Number of mutated TTCCG promoter hotspot sites per tumor, out of 291 in total as defined above, plotted against the whole-genome mutational burden across 221 melanomas. (**f**) TTCCG-related promoter hotspots arise preferably near highly expressed genes. 241 genes hosting 291 sites as defined above were considered. Expression levels were based on the median RPKM value across a subset of 38 TCGA melanomas with available RNA-seq. ND, not detected.

Of 51 recurrent promoter mutations (+/-500 bp from TSS) mutated in ≥ 12 tumors, 42 (82%) had a TTCCG element in the immediate (+/-10 bp) sequence context, rising to 86% after excluding the known *TERT* C228T and C250T promoter mutations (**Fig. 1b** and **Table S1**) ^1,2^. Most were within 200 bp upstream of a known TSS, as expected for functional ETS elements (**Fig. 1b-c**) ^17^. Among the few remaining sites, two (upstream of *AP3D1* and *TMEM102*) were instead flanked by TTCCT sequences likewise matching the ETS recognition motif (**Fig. 1b**) ^17^ and the numbers are thus conservative. The fraction TTCCG-related sites increased as a function of recurrence, from 291/550 promoter sites (53%) at *n* ≥ 5 to 7/8 (88%) at *n* ≥ 20, excluding the known *TERT* sites (**Fig. 1d**). For comparison, only 0.60% of C>T mutations in the dataset exhibited TTCCG patterns, underscoring their massive enrichment in recurrent positions.

As noted previously ^12^, there was a strong correlation between the number of mutated TTCCG hotspot sites and the total mutational burden in each tumor, compatible with these sites being passive passengers (Spearman’s *r* = 0.94, *P* = 1.5e-106; **Fig. 1e**). Also confirming earlier observations, the TTCCG-related promoter hotspots were found preferably near highly expressed genes, as expected under a model where interaction with an ETS TF rather than sequence-intrinsic properties are responsible for elevated mutation rates in these sites (**Fig. 1f**). Taken together, these analyses clearly demonstrate that ETS-related mutations account for nearly all highly recurrent non-coding hotspots genome-wide in melanoma, as well as hundreds of less recurrent sites not detectable in previous analyses based on smaller cohorts.

### TTCCG hotspots show elevated sensitivity to UV mutagenesis *in vitro* in the absence of repair

Recent studies have shown that NER, the main DNA repair pathway for UV damage, is attenuated in TF binding sites, leading to elevated somatic mutation rates^13,14^. While plausible as a mechanism for TTCCG mutation hotspots^11^, we recently showed that the hotspots appeared to be maintained in skin squamous cell carcinomas lacking global NER (*XPC* -/-). We also established that mutations can be easily induced in TTCCG hotspot sites in cell culture by UV light, thus recreating *in vitro* the process leading to recurrent mutations in tumors^12^. We decided to use the *RPL13A* -116 bp hotspot site, notably more frequently mutated (58/221 tumors) than both canonical *TERT* sites and on par with *BRAF* V600E at 60/221 (**Fig. 1ab**), as a model to further investigate a possible role for impaired NER.

To this end, we UV-exposed A375 cells with intact NER as well as fibroblasts with homozygous mutations in four key DNA repair components: *XPC*, required for global NER, *ERCC8* (*CSA*) and *ERCC6* (*CSB*), required for transcription coupled NER (TC-NER), and *XPA* which is required for lesion verification in both global and TC-NER (**Fig. S2**). Correct genetic identity and complete homozygosity for the mutant allele was confirmed by whole-genome sequencing of all four mutant cell lines (**Table S2**). Even limited UV exposure led to high cell mortality in the mutant cell lines, forcing us to limit the exposure to a single low dose of UVB (20 J/m^2^) during approximately two seconds, after which cells were assessed for *RPL13A* promoter mutations using error-corrected amplicon sequencing following recovery (**Fig. 2a**) ^18^. Between 7,332 and 13,774 error-corrected reads at ≥10x oversampling were obtained for each of 10 different libraries (**Fig. 2b**).

**Figure 2.**
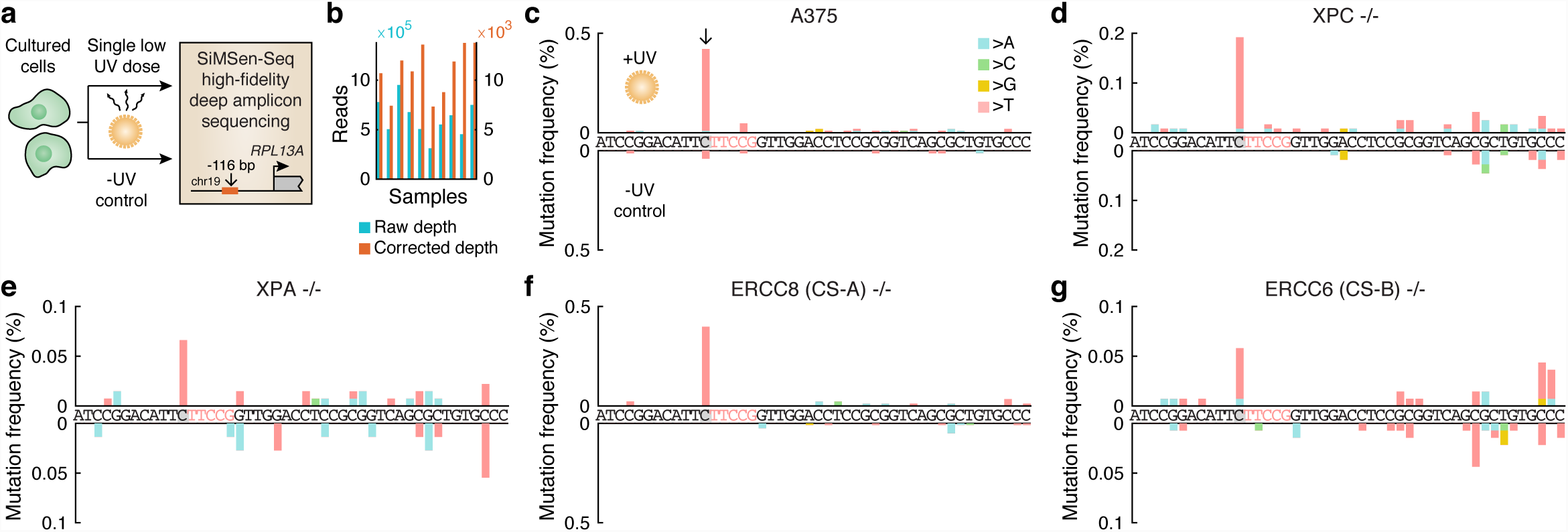
UV exposure of cultured cells induces mutations preferably at the *RPL13A* TTCCG hotspot site independently of repair. (**a**) Cultured human cells, either A375 melanoma cells or fibroblasts with NER deficiencies, were subjected to a single UVB dose (20 J/m^2^) during approx. 2 seconds. Following recovery, cellular DNA was subsequently assayed for subclonal mutation in the *RPL13A* -116 bp TTCCG promoter hotspot site (see **Fig. 1a** and **Fig. 1b**, top row) using SiMSen-Seq error-corrected amplicon sequencing^18^. Non-UV-treated sample were included as controls. (**b**) 10 samples were sequenced at 311k to 950k reads each, resulting in 7.3k to 13.8k error-corrected reads at ≥10x oversampling. (**c**) Subclonal mutations in a 46 bp amplicon window encompassing the *RPL13A* -116 bp hotspot in A375 melanoma cells. The hotspot site and TTCCG element are indicated in gray/red, respectively. Positive axis, UV-treated sample; negative axis, no UV control. (**d-g**) As panel **c** but showing results from *XPC* -/- (lacking global NER), *XPA* -/- (lacking global and transcription-coupled NER), *ERCC8* -/- and *ERCC6* -/- (lacking transcription-coupled NER) mutant fibroblasts.

Strikingly, even at this miniscule dose, subclonal somatic mutations appeared preferably at the known hotspot site in A375 cells (**Fig. 2c**) as well as in all of the mutant cell lines (**Fig. 2e-g**), despite abundant possibilities for UV lesion formation in flanking assayed positions. As expected, absolute mutation frequencies were low, less than 0.5% in all samples, bringing us close to the detection limit in some samples as indicated by noise in the untreated controls (**Fig. 2d,e,g**). In combination with earlier data from *XPC* -/- tumors lacking global NER ^12^ and the fact that the mutations are almost exclusively positioned upstream of TSSs where TC-NER should not be active (**Fig. 1bc**), these results argue strongly against impaired NER as a mechanism for TTCCG hotspot formation.

### High-resolution mapping of CPDs across the human genome

It was established decades ago that DNA conformational changes induced by interactions with proteins can alter conditions for UV damage formation^19,20^, which prompted us to investigate whether ETS-related promoter hotspots may arise due to locally favorable conditions for UV lesion formation rather than inhibited repair. For this, we adapted a protocol first established in yeast using IonTorrent sequencing^21^ to the Illumina platform (**Fig. 3a**), to generate a genome-wide map of CPDs in A375 human melanoma cells immediately following UV exposure, before DNA repair processes have had a chance to act.

**Figure 3.**
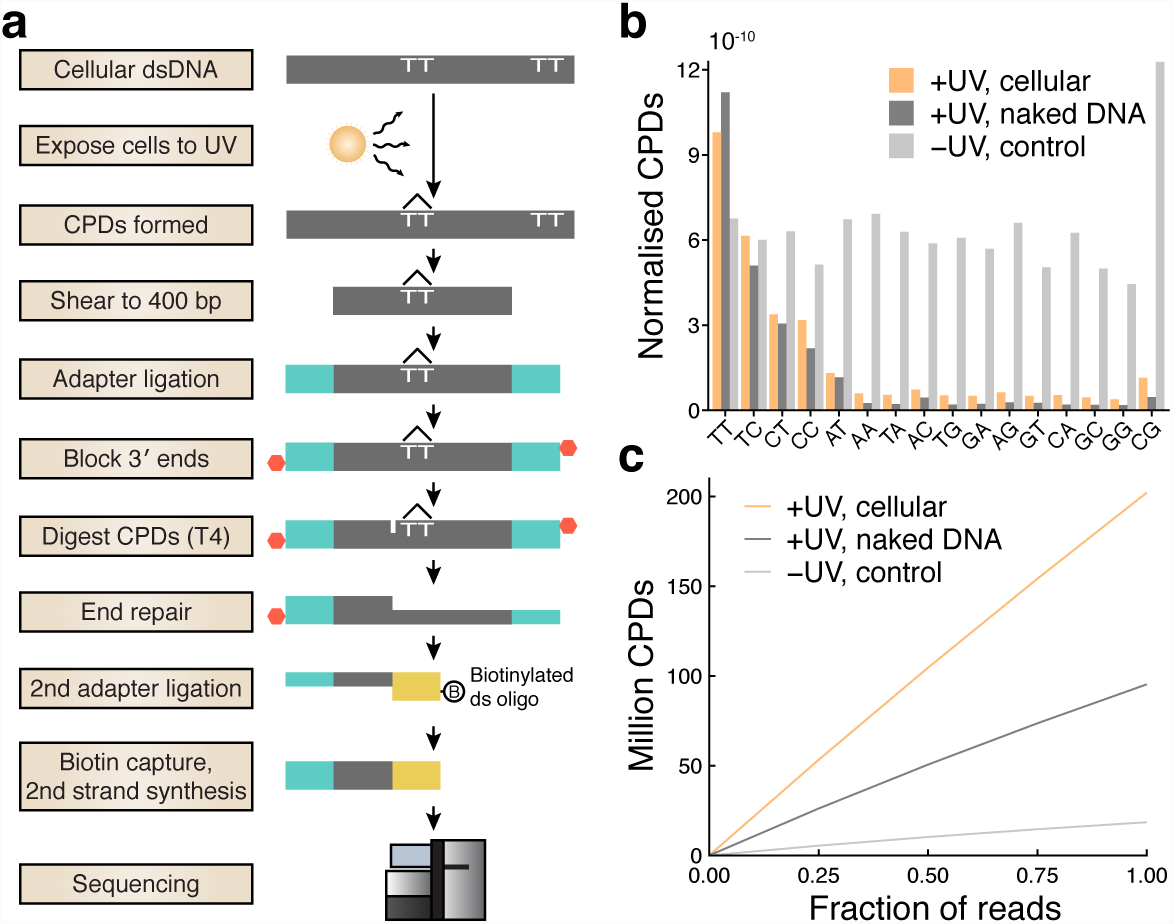
High-coverage mapping of UV-induced cyclobutane pyrimidine dimers across the human genome. (**a**) Schematic of the experimental protocol. (**b**) Distribution of dinucleotides at which CPDs were detected, showing an expected preference for dipyrimidines. Counts from cellular, naked (acellular) and no-UV-control samples were normalized with respect to genomic dinucleotide counts as well as sequencing depth. (**c**) The number of detected CPDs in each library following removal of PCR duplicates shown at full depth, as well as based on subsampled data (25, 50 and 75%) to simulate lower sequencing depth.

CPDs were preferably detected at TT, TC, CT and CC dinucleotides as expected (**Fig. 3b**) and the number of detected CPDs after removal of PCR duplicates scaled nearly linearly with simulated sequencing depth, indicating favorable random representation of CPDs (**Fig. 3c**). A total of 202.1 million CPDs were mapped to dipyrimidines throughout the genome (**Fig. 3c**), constituting the highest resolution CPD map to date to our knowledge. Additionally, 95.3 million CPDs were mapped in UV-treated naked (acellular) A375 DNA lacking interacting proteins, while a non-UV-treated control, which expectedly yielded limited material, produced 18.5 million CPDs (**Fig. 3c**).

### CPD formation spikes at TTCCG-related promoter mutation hotspots

We next investigated CPD formation patterns at TTCCG mutation hotspots positions identified above in melanoma (**Fig. 1a-b**). 291 recurrently mutated (*n* ≥ 5/221 melanomas) TTCCG promoter sites (+/-500 bp from TSS) were aligned centered on the mutated base such that CPD density in these regions could be determined. This revealed a striking peak in CPD formation that coincided with the hotspots, which was largely absent in naked DNA lacking bound proteins or in non-UV control DNA (**Fig. 4a**). Additionally, more recurrently mutated sites showed a stronger CPD signal, compatible with increased CPD formation being the key mechanism (**Fig. 4b**).

**Figure 4.**
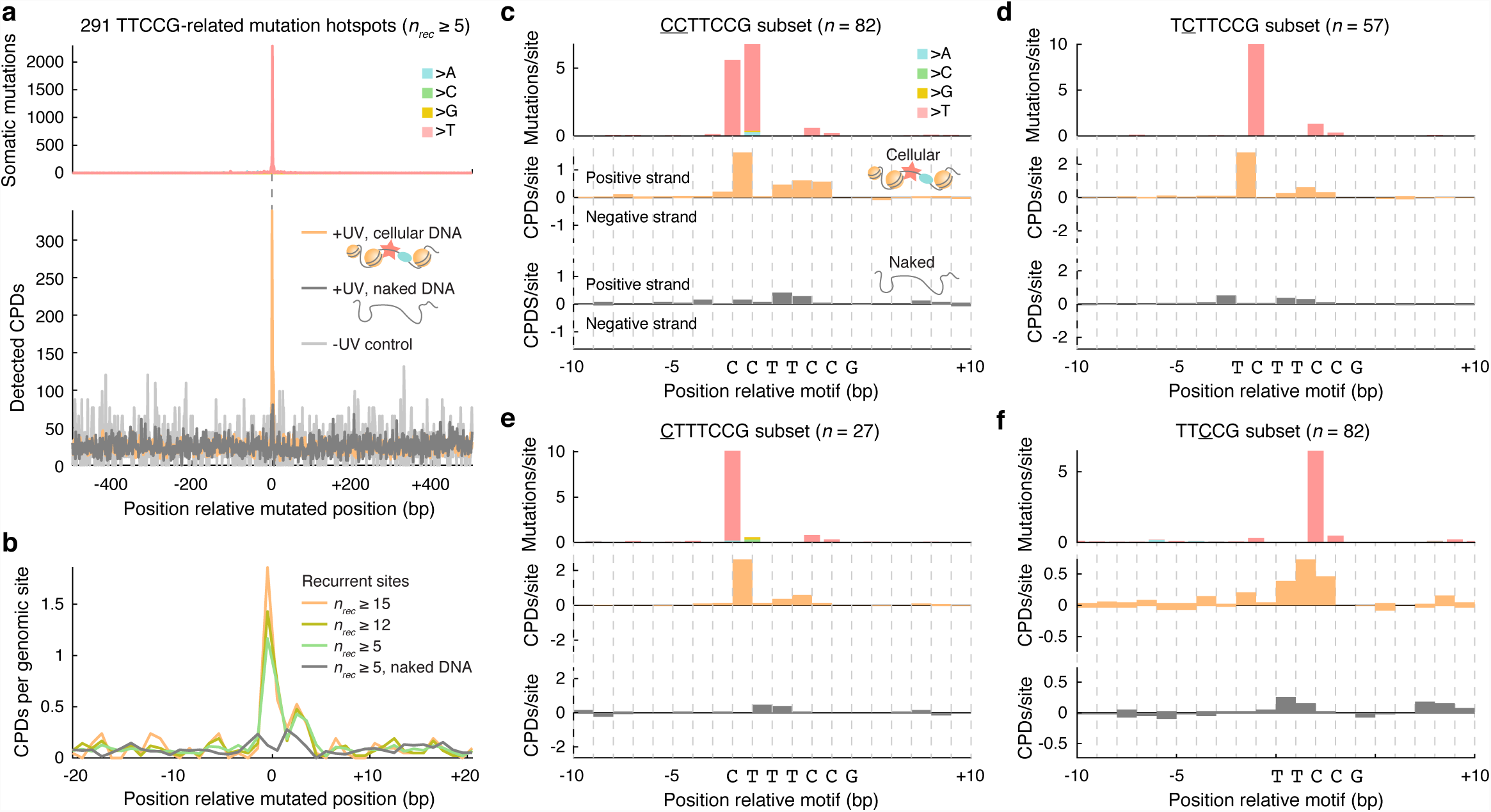
CPD formation spikes at TTCCG-related promoter mutation hotspots. (**a**) 291 recurrently mutated (*n* ≥ 5/221 melanomas) genomic promoter sites (+/-500 bp from nearest TSS), as defined and illustrated in **Fig 1a-b,** were aligned centered on the mutated base (in each case considering the pyrimidine-containing strand, i.e. C, for the mutated base in the reference genome). The top and bottom panels show mutation and CPD formation density, respectively, in a +/- 500 bp window centered on the mutated base. Naked DNA (dark grey) and no-UV control (light grey) whole-genome CPD counts were normalized to be comparable to the cellular DNA data (orange). (**b**) Close-up view (+/- 20 bp) showing CPD density for different subsets of the 291 sites, defined by the degree of mutation recurrence, revealing that more prominent melanoma mutation hotspots show stronger CPD formation signals. (**c-f**) Detailed view of CPD formation patterns in TTCCG promoter mutation hotspot sites after subcategorization into four main groups based on sequence and mutated position (mutated base indicated by underscore, with <u>CC</u>TTCCG sites typically showing recurrent mutations at both 5’ cytosines). Mutated genomic regions were aligned centered at the start of the motif while removing redundant (non-unique) genomic loci. Mutation and CPD formation frequencies were normalized by the number of hotspot sites in each alignment, following depth-normalization as described in panel **a**. CPD frequencies are shown separately for the positive and negative strands, for both cellular (orange) and naked (grey) DNA.

For a more detailed understanding, we subcategorized the 291 melanoma ETS hotspot sites into four main groups based on sequence and mutated position. The strongest mutation hotspots, such as *RPL13A* and *DPH3*, typically occurred at cytosines one of two bases upstream of the TTCCG element (**Fig. 1b** and **Table S1**), which notably is outside of the core motif and therefore not expected to disrupt binding ^17^. In <u>CC</u>TTCCG sites (*n* = 82 unique loci), recurrent C>T transitions would typically appear at both 5′ cytosines (underscored) independently or, less frequently, as CC>TT double nucleotide substitutions. Aggregated CPD density overs these sites, centered on the motif, revealed a strong peak bridging these two bases, which notably was absent in naked DNA (**Fig. 4c**). Thus, when the TF site is occupied, CPDs form efficiently between the two pyrimidines, leading to C>T mutations at either base although with a preference for the second position, in agreement with established models for UV mutagenesis ^22^. The same pattern of strongly elevated CPD formation in cellular, but not naked, DNA was observed between the same positions in T<u>C</u>TTCCG and C<u>T</u>TTCCG sites (*n* = 57 and 27, respectively), with C>T mutations expectedly forming only at the first or second pyrimidine depending on the position of the cytosine (**Fig. 4d-e**).

Many of the less recurrent bases in melanoma were often found at the first middle cytosine of a TT<u>C</u>CG motif (**Table S1**). Interestingly, a large fraction of these sites lacked a dipyrimidine at the two key positions identified above thus prohibiting CPD formation there, with ACTT<u>C</u>CG being the most common pattern (44/82 sites), which indeed matches the *in vivo* ETS consensus sequence ^17^. Compatible with the mutation data, the strongest CPD peak was observed at the middle T<u>C</u> dinucleotide, and in agreement with the lower mutation recurrence, this signal was weaker compared to the other site types (**Fig. 4f**). Of note, elevated CPD formation between these bases could also be clearly seen in the other site categories (**Fig. 4c-e**). Taken together, these analyses based on genome-wide CPD mapping provide strong evidence that locally elevated CPD formation efficacy underlies the formation of mutation hotspots at ETS binding sites.

### Overall elevated mutation rate in regulatory regions is not due to increased CPD formation

Earlier studies have described a general increase in mutation rate in promoter regions, attributed to reduced NER activity at sites of TF binding ^13,14^. To investigate a possible contribution from increased CPD formation to this pattern, we first determined the overall mutation rate near TSSs, which confirmed a sharp increase in upstream regions that coincided with reduced NER as determined by XR-Seq (**Fig. 5a-b**) ^23^. However, aggregated over these regions, CPDs were found to form at near-expected frequencies (**Fig. 5c**). While individual ETS-related mutation hotspots arise due to elevated CPD formation, impaired NER thus appears to be the main contributor to a general increase in mutation burden in promoters.

**Figure 5.**
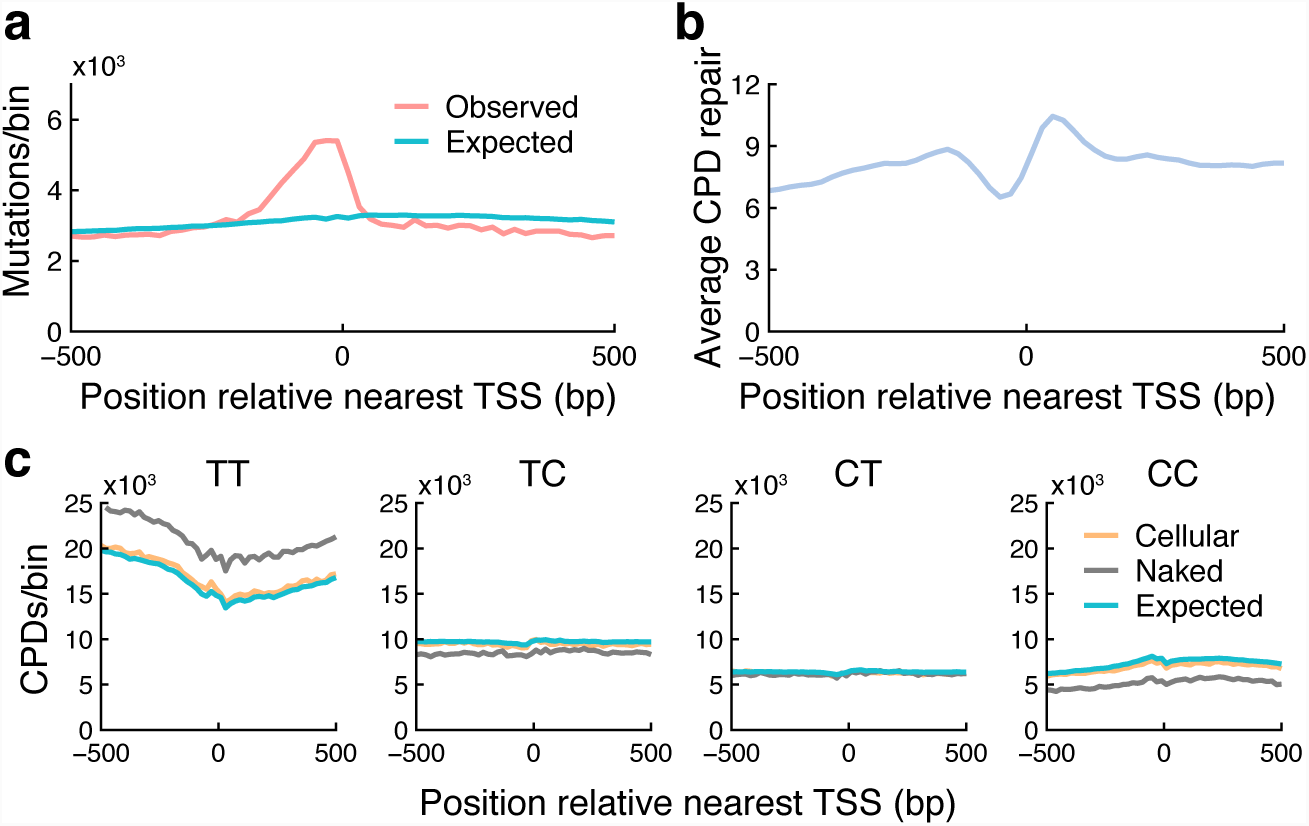
Overall elevated mutation rate in regulatory regions is not due to increased CPD formation. (**a**) Somatic C>T mutation density around annotated TSSs, per 20 bp genomic bin, aggregated over 33,456 coding genes and lncRNAs in the GENCODE v19 annotation. Expected mutation counts were determined by generating an equal number of mutations using observed trinucleotide mutational signatures in the analyzed samples. (**b**) Average NER activity as determined by XR-Seq^23^ in TSS regions. (**c**) Observed CPD counts in TSS regions (20 bp bins) in cellular and naked DNA, presented per CPD-forming dinucleotide. Naked DNA counts were normalized to be comparable to the cellular DNA data. Expected counts were determined based on dinucleotide counts in the analyzed regions.

## DISCUSSION

Proper analysis of recurrent non-coding mutations requires an understanding of how mutations arise and distribute across the genome in the absence of selective pressures. Here, we provide a mechanistic explanation for the passive emergence of recurrent mutations at TTCCG/ETS sites in tumors in response to UV light, and also demonstrate their massive impact on the mutational landscape of melanoma using a large whole genome cohort.

Mutations at -116 bp in the *RPL13A* promoter were used here as a model to study mutation formation at ETS hotspot sites *in vitro* in repair-deficient cell lines, which ruled out an important role for inhibited repair. Of note, this site is more recurrently mutated than the individual *TERT* C228T/C250T sites and nearly as frequent as chr7:140453136 mutations (hg19) pertaining to *BRAF* V600E, thus representing the second most common mutation in melanoma and likely other skin cancers. Notably, mutations were detectable at this site in cultured cells following a UVB dose of 20 J/m^2^ UVB, equivalent to about 1/200th of the monthly absorbed UVB dose in July in Northern Europe^24^. This underscores the extreme UV sensitivity of ETS hotspots and explains their high recurrence in tumors.

Genome-wide mapping of CPDs revealed that TTCCG-related mutation hotspots exhibit highly efficient CPD formation at the two bases immediately 5’ of the core TTCC ETS motif. The effect was lost in naked acellular DNA, showing that structural conditions for elevated CPD formation are induced when the TF binding site is in its protein-bound state. Interestingly, most functional ETS sites are expected to lack pyrimidines in the two key positions ^17^ thus prohibiting pyrimidine dimer formation, and conditions for forming a strong mutation hotspot are thus only met in a subset of sites with CC, TC or CT preceding the TTCCG element. Additionally, CPDs form at lower but still elevated frequency at the middle T<u>TC</u>CG bases, consistent with weaker recurrence for mutations in these positions. CPD and cancer genomic data are thus in strong agreement, providing a credible mechanism for the formation of ETS-related mutations hotspots in UV-exposed cancers.

As demonstrated here, frequent mutations at ETS-site hotspots are expected for purely biochemical reasons in UV-exposed cancers. Consequently, several observations are compatible with passenger roles for these mutations: The most recurrent sites arise at cytosines outside of the core TTCC ETS recognition element ^17^ where they are not expected to disrupt ETS binding. While mutations in the middle of the motif, common among the less frequent hotspots, should disrupt binding, ETS factors tend to be oncogenes that are activated in cancer ^25^, and it can be noted that *TERT* promoter mutations instead enable ETS binding through formation of TTCC elements ^1,2^. The mutations tend to arise near highly expressed housekeeping genes rather than cancer-related genes. Moreover, as would be expected in the absence of selection and in contrast to known driver mutations^12^, the number of mutated ETS-sites in a tumor is strongly determined by mutational burden.

Our results complement a recent study by Mao, Brown ^26^ et al., which was published during the preparation of this manuscript. This study likewise determined CPD formation patterns in ETS binding sites, obtaining results that are in full agreement with ours, and also proposed a structural basis for increased CPD formation in the ETS-DNA complex based on available crystallographic data. While inhibited NER appears to explain the majority of the increased mutation burden in regulatory DNA in UV-exposed cancers, it can thus be concluded with confidence that the strongest individual mutation hotspots arise instead due to base-specific elevations in CPD formation efficacy at ETS TF binding sites.

## METHODS

### Whole genome mutation analyses

Whole genome somatic mutation calls from the Australian Melanoma Genome Project (AMGP) cohort ^15^ were downloaded from the International Cancer Genome Consortium’s (ICGC) database ^27^. These samples were pooled with whole genome mutation calls from The Cancer Genome Atlas (TCGA) melanoma cohort ^16^ called as described previously ^5^. Population variants (dbSNP v138) and duplicate samples from the same patient were removed, resulting in a total of 221 tumors.

Gene annotations from GENCODE^28^ v19 were used to define TSS positions, encompassing 20,149 and 13,307 uniquely mapped coding genes and lncRNAs, respectively, considering the 5’-most annotated transcripts while disregarding non-coding isoforms for coding genes. Processed RNA-seq data was derived from Ashouri, Sayin ^29^.

### Culture and UV treatment of repair-deficient fibroblasts

XP12, GM16094, GM16095 and GM15893 cells were a kind gift from Dr. Isabella Muyleart, University of Gothenburg. Cells were grown in DMEM + 10% FCS + Penicillin/streptomycin (GIBCO). Cells were subjected to a single low dose UVB (20 J/m^2^) and left to recover. DNA was extracted with Blood Mini kit (Qiagen).

### Ultrasensitive mutation analysis

To detect and quantify mutations we applied SiMSen-Seq (Simple, Multiplexed, PCR-based barcoding of DNA for Sensitive mutation detection using Sequencing) as described in Fredriksson et al 2017. Sequencing was performed on an Illumina MiniSeq instrument in 150 bp single-end mode. Raw FastQ files were subsequently processed as described using Debarcer Version 0.3.1 (https://github.com/oicr-gsi/debarcer/tree/master-old). For each amplicon, sequence reads containing the barcode were grouped into barcode families. Barcode families with at least 10 reads, where all of the reads were identical (or ≥ 90% for families with >20 reads), were required to compute consensus reads. FastQ files were deposited in the Sequence Read Archive under BioProject ID PRJNA487997.

### Genome-wide mapping of cyclobutane pyrimidine dimers

A375 cells were grown in DMEM + 10% FCS + Penicillin/streptomycin (GIBCO) and were treated with 1000 J/m^2^ UVC following DNA extraction and DNA from untreated cells was isolated as a control, both in duplicates. Additionally, naked DNA from untreated cells was irradiated with the same dose, to provide an acellular DNA control sample. DNA was extracted with the Blood mini kit (Qiagen). Purified DNA (12 ug) was sheared to 400bp with a Covaris S220 in microtubes using the standard 400 bp shearing protocol. CPD-seq was modified from Mao, Smerdon ^21^ to adapt it to Illumina sequencing methods using primers described previously in Clausen, Lujan ^30^ (**Table S3**). Briefly, sheared DNA was size selected with SPRI select beads (Life Technologies) and the purified product (approx. 4 ug) subjected to NEBNext end repair and NEBNext dA-tailing modules (NEB). ARC141/142 was then ligated to the sheared and repaired ends O/N with NEBNext Quick Ligation module. DNA was purified with CleanPCR beads and treated with Terminal Transferase (TdT, NEB) and dideoxy ATP (Roche) for 2h at 37 degrees. DNA was purified and incubated with 30 units T4 endonuclease V (NEB) at 37 degrees for 2 h, followed by purification and treatment with APE1 (NEB) at 37 degrees for 1.5 hr. DNA was purified and treated with rSAP (NEB) 37 degrees 1 hr followed by deactivation at 65 degrees for 15 minutes. DNA was purified, denatured at 95 degrees for 5 min, cooled on ice and ligated with the biotin-tagged “ARC double” overnight at 16 degrees with NEBNext quick ligation module. DNA fragments with the biotin tag were captured with Streptavidin Dynabeads (Invitrogen) and the DNA strand without the biotin label was released with 0.15 M NaOH. This single stranded DNA was used as the template to synthesise double stranded products using ARC153. The now double stranded library was purified and amplified with ARC140 and ARC78-82 to add Illumina barcodes and indexes. Two cellular UV-treated, two no-UV controls and one naked DNA control library was prepared, for a total of five libraries. The libraries were pooled with equal volumes of each the libraries and sequenced using a NextSeq High Output kit (Illumina).

### CPD bioinformatics

FastQ files were aligned pairwise with Bowtie 2 version 2.3.1 ^31^ to hg19, using standard parameters. For the -UV control and +UV cellular DNA samples, replicates were merged with Picard MergeSamFiles version 2.18.7 (http://broadinstitute.github.io/picard). Duplicate reads were marked with Picard MarkDuplicates version 2.18.7^32^ with the parameter VALIDATION_STRINGENCY=LENIENT. Further analysis was performed in R with Bioconductor ^33^, where CPD positions were extracted as the two bases upstream and on the opposite strand of the first mate in each read pair, removing those that mapped outside of the chromosome boundaries. Only biologically possible CPDs detected at dypyrimidines sites were considered in the CPD counts and downstream analyses. Data from duplicate libraries were pooled to achieve higher coverage, since downstream results were in close agreement when considering these libraries individually. To simulate lower coverage libraries, the bam files were subsampled with samtools view version 0.1.19-44428cd ^34^ with the parameter -s at 0.25, 0.5 or 0.75, and the subsequent bam files were reanalyzed as described above.

For analyses of CPD formation patterns, C>T mutations and repair activity around TSSs, these regions were divided into 20 bp bins in which CPD counts or overlapping XR-seq reads were determined. XR-seq data from wild-type NHF1 skin fibroblasts was obtained from ^35^, and consisted of normalized read counts in 25 bp strand-specific bins. Background frequencies of dinucleotides and trinucleotides in hg19 were counted with EMBOSS’s fuzznuc ^36^, using the parameters -auto T -complement T. Expected mutations were calculated by randomly introducing the same number of mutations as observed in the window based on observed probabilities for C>T mutations at different trinucleotides estimated from the complete mutation dataset. Expected CPDs were calculated in the same way, maintaining the number of CPDs in the observed data, but based instead on genomic dinucleotide counts.

## ACKNOWLEDGEMENTS

The results published here are in whole or part based upon data generated by The Cancer Genome Atlas pilot project established by the NCI and NHGRI, as well as ICGC. Information about TCGA and the investigators and institutions who constitute the TCGA research network can be found at “http://cancergenome.nih.gov”. We are most grateful to the patients, investigators, clinicians, technical personnel, and funding bodies who contributed to TCGA and ICGC, thereby making this study possible. This work was supported by grants from the Knut and Alice Wallenberg Foundation (E.L, A.S), the Swedish Foundation for Strategic Research (E.L), the Wenner-Gren Foundation (E.L), the Swedish Medical Research Council (E.L., A.S), the Swedish Cancer Society (E.L, A.S), the Swedish Childhood Cancer Foundation (A.S), Sahlgrenska Academy (ALF) at University of Gothenburg (A.S.), the Åke Wiberg foundation (E.L), and the Lars Erik Lundberg Foundation for Research and Education (E.L.). The computations were in part performed on resources provided by SNIC through Uppsala Multidisciplinary Center for Advanced Computational Science (UPPMAX) under project b2012108.

## AUTHOR CONTRIBUTIONS

M.B, E.L, K.E, S.F, M.L and J.VdE performed bioinformatics analyses; K.E, E.L, A.C and A.S designed experiments; K.E. and S.F. performed experiments; E.L wrote the paper with contributions from K.E and M.B; E.L. conceived the study.

## SUPPLEMENTARY FIGURES

**Figure S1.**
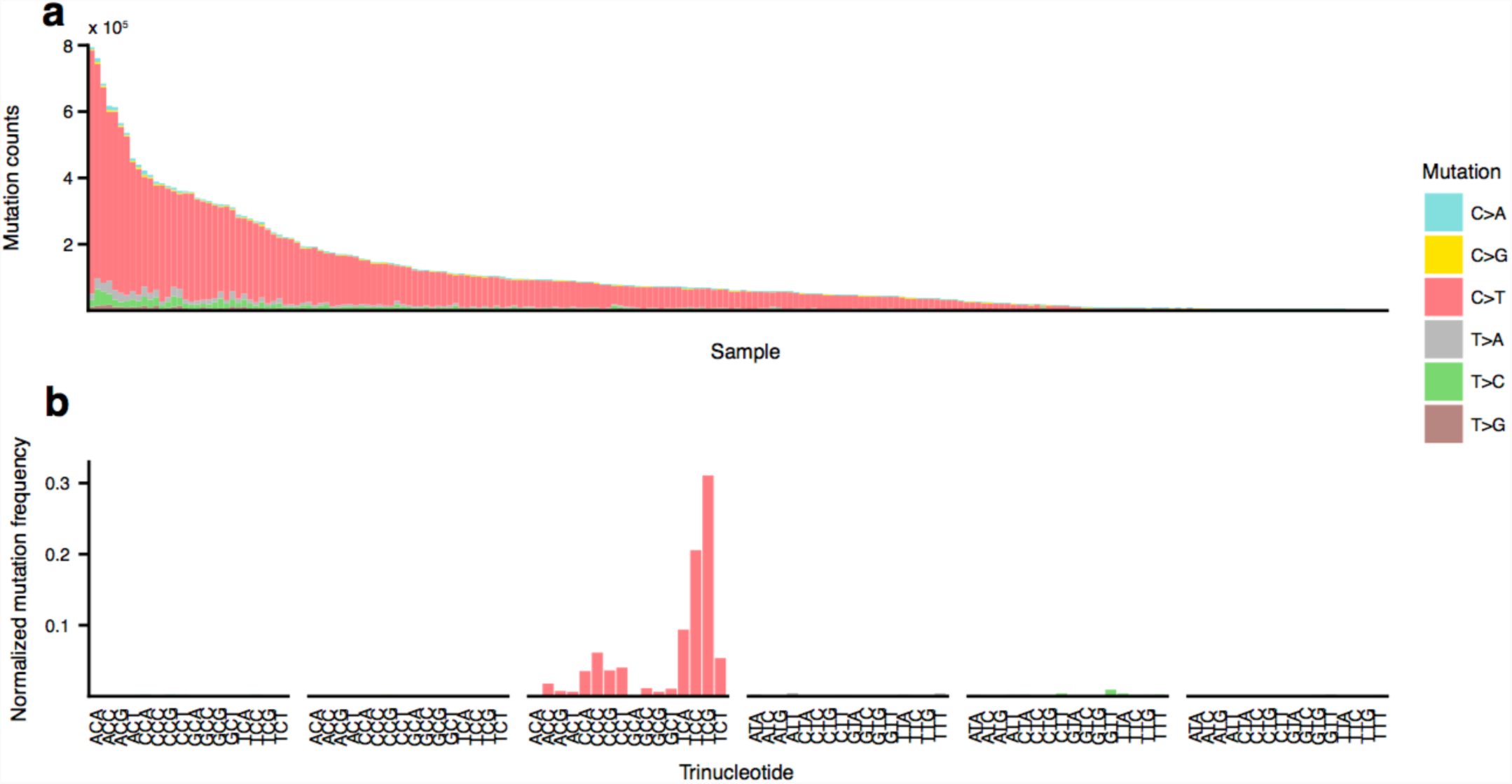
Mutational burden and overall mutational signature for 221 melanomas. (**a**) Number of mutations in each sample, color-coded for pyrimidine-based nucleotide substitution. (**b**) Mutation frequency of each substitution type in different trinucleotide contexts, normalized for genomic trinucleotide background frequencies.

**Figure S2.**
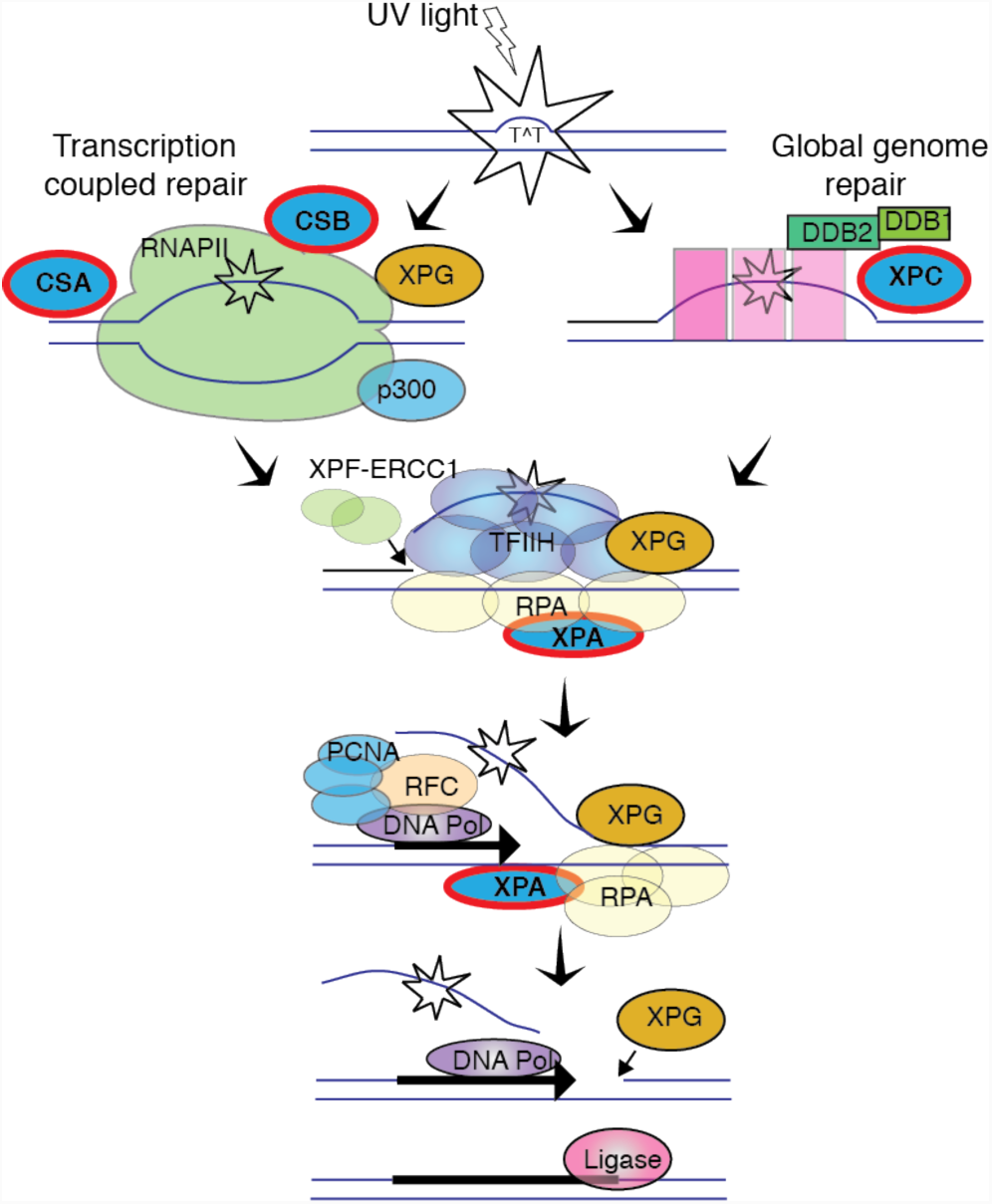
The nucleotide excision repair (NER) pathway, with mutated genes in the four repair-deficient cell lines (Table S1) highlighted in red. Adapted from Hanawalt and Spivak ^1^.

## SUPPLEMENTARY TABLES

**Table S2.** Cell lines with DNA repair deficiencies and their verified homozygous mutations

| Cell line | Gene | hg19 position | Protein substitution | Nucleotide substitution |
| --- | --- | --- | --- | --- |
| XP12RO | *XPA* | Chr9:100447259 | p.R207X | c.619 C>T |
| GM16094 | *ERCC8 (CSA)* | Chr5:60200621 | p.Y145X | c.435 T>G |
| GM16095 | *ERCC6 (CSB)* | Chr10:50732467 | p.K337X | c.1009 A>T |
| GM15983 | *XPC* | Chr3:14199740 | p.V548delTG | c.1643_1644delTG |

**Table S3.**
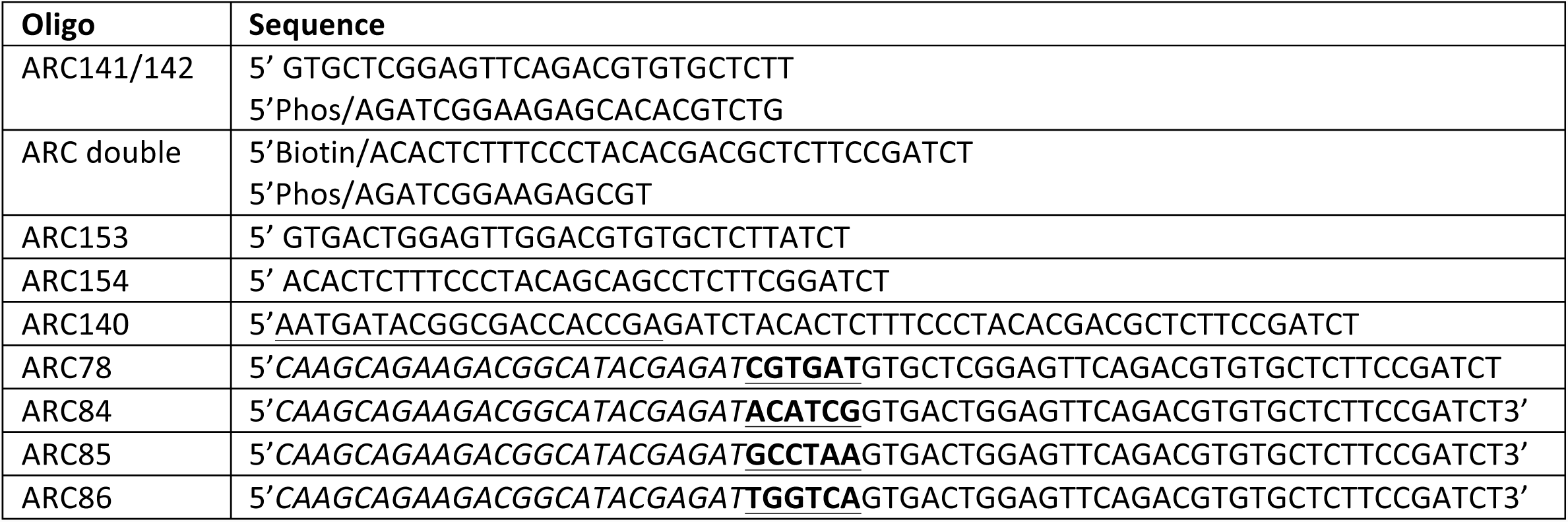
Oligonucleotide sequences for CPD-seq. Illumina <u>P5</u> and *P7* adapters are indicated underlined and italicized respectively, and **<u>indexes</u>** are shown in bold and underline. Oligo 5’ modifications are also indicated. All oligos were from Integrated DNA technologies (Coralville, IA)

